# Telomere-to-Telomere and Haplotype-Phased Genome Assemblies of the Heterozygous Octoploid ‘Florida Brilliance’ Strawberry (*Fragaria × ananassa*)

**DOI:** 10.1101/2022.10.05.509768

**Authors:** Hyeondae Han, Christopher R Barbey, Zhen Fan, Sujeet Verma, Vance M. Whitaker, Seonghee Lee

**Author notes:** Correspondence: Seonghee Lee, IFAS Gulf Coast Research and Education Center, 14625 CR 672, Wimauma, FL, 33598, USA.

## Abstract

The available haplotype-resolved allo-octoploid strawberry (*Fragaria* × *ananassa* Duch.) (2*n* = 8*x* = 56) genomes were assembled with the trio-binning pipeline, supplied with parental short-reads. We report here a high-quality, haplotype-phased genome assembly of a short-day cultivar, ‘Florida Brilliance’ (FaFB2) without the use of parental sequences. Using Pacific Biosciences (PacBio) long reads and high-throughput chromatic capture (Hi-C) data, we completed telomere-to-telomere phased genome assemblies of both haplotypes. The N50 continuity of the two haploid assemblies were 23.7 Mb and 26.6 Mb before scaffolding and gap-filling. All 56 pseudochromosomes from the phased-1 and phased-2 assemblies contained putative telomere sequences at the 5’ and/or 3’ ends. A high level of collinearity between the haplotypes was confirmed by high-density genetic linkage mapping with 10,269 SNPs, and a high level of collinearity with the ‘Royal Royce’ FaRR1 reference genome was observed. Genome completeness was further confirmed by consensus quality. The LTR assembly Index score for entire genome assembly was 19.72. Moreover, the BUSCO analysis detected over 99% of conserved genes in the combined phased-1 and phased-2 assembly. Both haploid assemblies were annotated using Iso-Seq data from six different ‘Florida Brilliance’ tissues and RNA-Seq data representing various *F*. × *ananassa* tissues from the NCBI sequence read archive, resulting in a total of 104,099 genes. This telomere-to-telomere reference genome of ‘Florida Brilliance’ will advance our knowledge of strawberry genome evolution and gene functions, and facilitate the development of new breeding tools and approaches.

## INTRODUCTION

Cultivated strawberry (*Fragaria* × *ananassa* Duch.) is an allo-octoploid species (2n = 8x = 56) originating from the spontaneous interspecific hybridization of two wild octoploid species *F*. *chiloensis* and *F*. *virginiana* (1). It is believed that the octoploid progenitors of cultivated strawberry were generated by successive stages of polyploidization of four diploid progenitor species more than one million years (2). The consensus diploid progenitor species are identified as *F*. *vesca* and *F*. *iinumae* (2, 3).

De novo genome assembly is the most ideal and comprehensive and unbiased method to assemble DNA sequences. The first chromosome-scale genome assembly for octoploid strawberry of ‘Camarosa’ (2) was constructed with Illumina Solexa short reads (<1 kb) with a low rate of error (< 0.1%) and Pacific Biosciences (PacBio) long reads with a higher rate of error (<10%). The availability of the reference sequences contributed not only to our knowledge of the evolutionary history of octoploid strawberry, but also to the characterization of genetic loci for agronomic traits such as disease resistance and fruit quality (4). However, the assembly had limitations for wider use due to a lack of haplotype-phasing, phase-switching, assembly gaps, and scaffolding and local assembly errors that were expected with the available technology in a heterozygous octoploid.

A great challenge for haplotype-phased genome assembly is base-calling accuracy. Recently, the availability of high-fidelity (HiFi) reads developed by PacBio has become the foundation for phased genome assemblies. The first haplotype-phased genome assembly of cultivated strawberry was of the day-neutral octoploid strawberry variety ‘Royal Royce’, using trio-binning with parental short-read sequencing data (5). Sometimes, however, parental information is lacking. Recently, a new algorithm combining PacBio HiFi reads and Hi-C chromatin interaction data (6) generated fully haplotype-phased genome assemblies in a human genome study without parental information (7).

In the present study, we developed a genome assembly from a short-day strawberry variety ‘Florida Brilliance’, which is utilized in annual growing systems for winter and early spring production. ‘Florida Brilliance’ was released from the University of Florida strawberry breeding program and is currently the leading variety grown in Florida (8). Here, we present a telomere-to-telomere octoploid strawberry genome consisting of two haploid assemblies (phased-1 and phased-2), enabled by a single-sample method using HiFi and Hi-C data without parental information.

## RESULTS

### Octoploid Strawberry Genomic and Transcriptomic Datasets

The details of the sequencing read data utilized are provided in Table 1. With five single-molecular real-time cells in PacBio Sequel II platform, 144.1 Gb of sequence were generated in 9.1M reads. The average read length was 15,834.8 bp. The Hi-C data was generated on Novaseq6000 and contained 86.5 Gb of sequences in 286 M paired-end reads with an average length of 151 bp. In total, 6.56 Gb of ‘Florida Brilliance’ Iso-Seq data with an average length of 2,558 bp were generated. Among six tissues, when restricted to HiFi (QV≥20), 45,577 to 56,707 reads with an average length of about 2,500 bp were obtained (Supplementary Table 1). In total, 12.1 Gb of ‘Royal Royce’ Iso-Seq sequence data (5) and 511 Gb of short read Read-seq were used as input for transcriptome assembly and gene annotation.

**Table 1.**
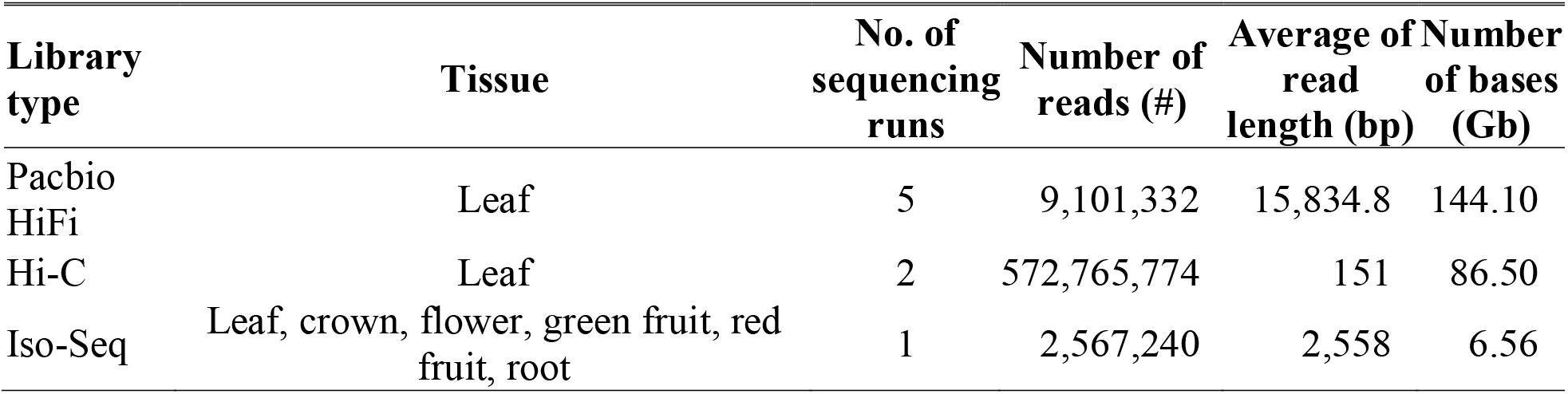
Libraries sequenced and used in assembly with accession numbers.

### Phased Octoploid Genome Assembly

The octoploid strawberry genome of ‘Florida Brilliance’ was assembled using HiFi and Hi-C reads. A smudgeplot based on K-mer analysis using HiFi reads was consistent with ‘Florida Brilliance’ having a genome structure of ‘AAAAAABB’ (Figure 1). The phased-1 assembly contained 3,716 contigs with an N50 of 23.7 Mb, and the phased-2 assembly contained 1,226 contigs with an N50 of 26.7 Mb (Table 2). Fifteen contigs accounted for 50% of the phased-2 assembly, indicating that one contig corresponds to a chromosome (Table 2). In addition, the largest contig sizes in the phased-1 and phased-2 genome assemblies were over 36 Mb (Table 2). Before scaffolding, the Benchmarking Universal Single-Copy Orthologs (BUSCO) scores were 99.2% in the phased-1 assembly and 99.1% in the phased-2 assembly, indicating the high-quality of the initial assembly (Table 2). Comparison of the full assembly to whole genome sequencing HiFi reads of FaFB2 using Merqury showed very high base accuracy (QV>69.8), indicating 99.99999% of HiFi reads were detected on the combined phased −1 and −2 contigs (Table 2). The final assembly contained 99.1% complete gene models with a majority (96.6%) of the duplicated complete gene models in both phased-1 and phased-2 genome assemblies (Table 2).

**Table 2.**
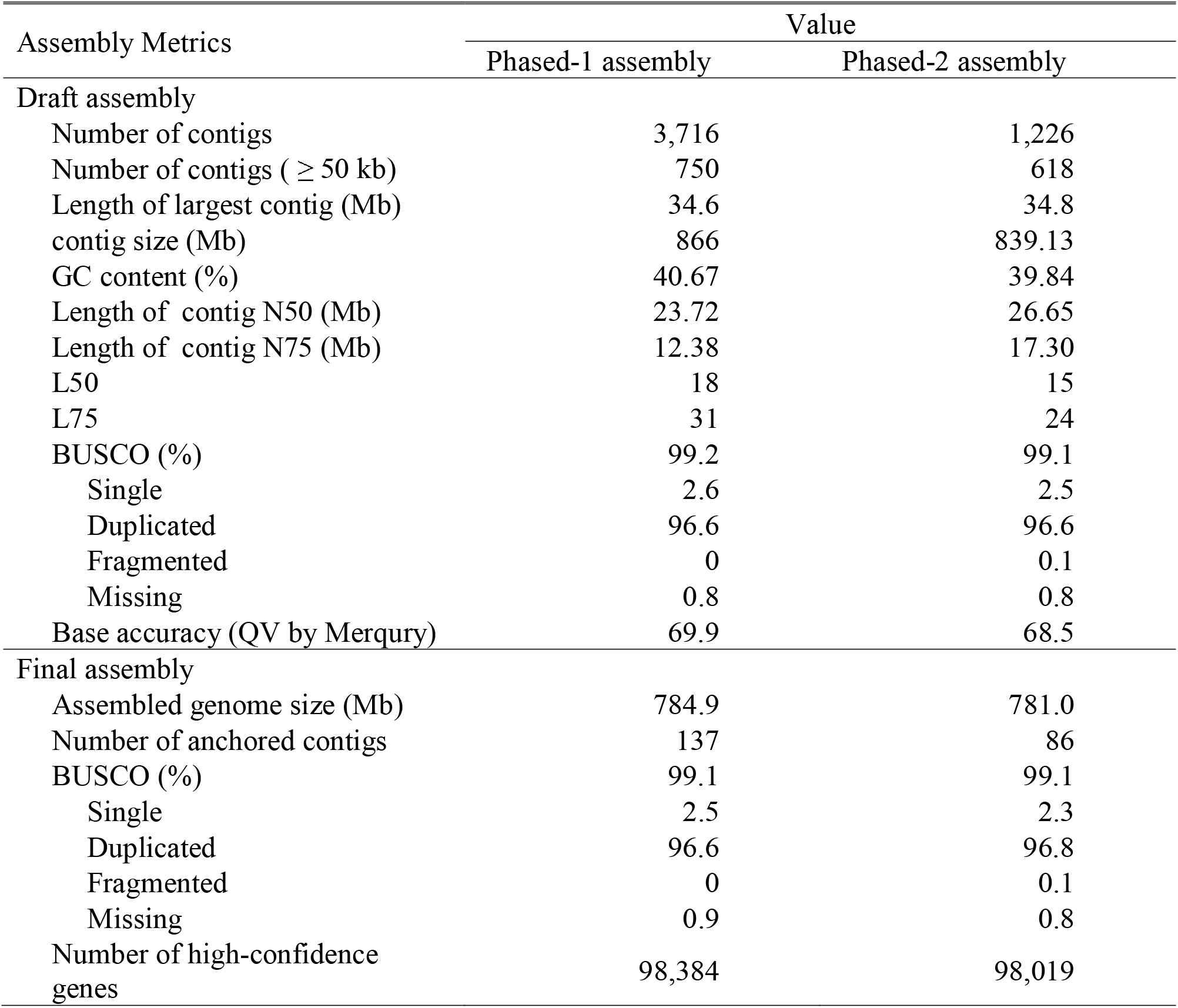
Statistics of the ‘Florida Brilliance’ genome assembly and annotation.

**Figure 1.**
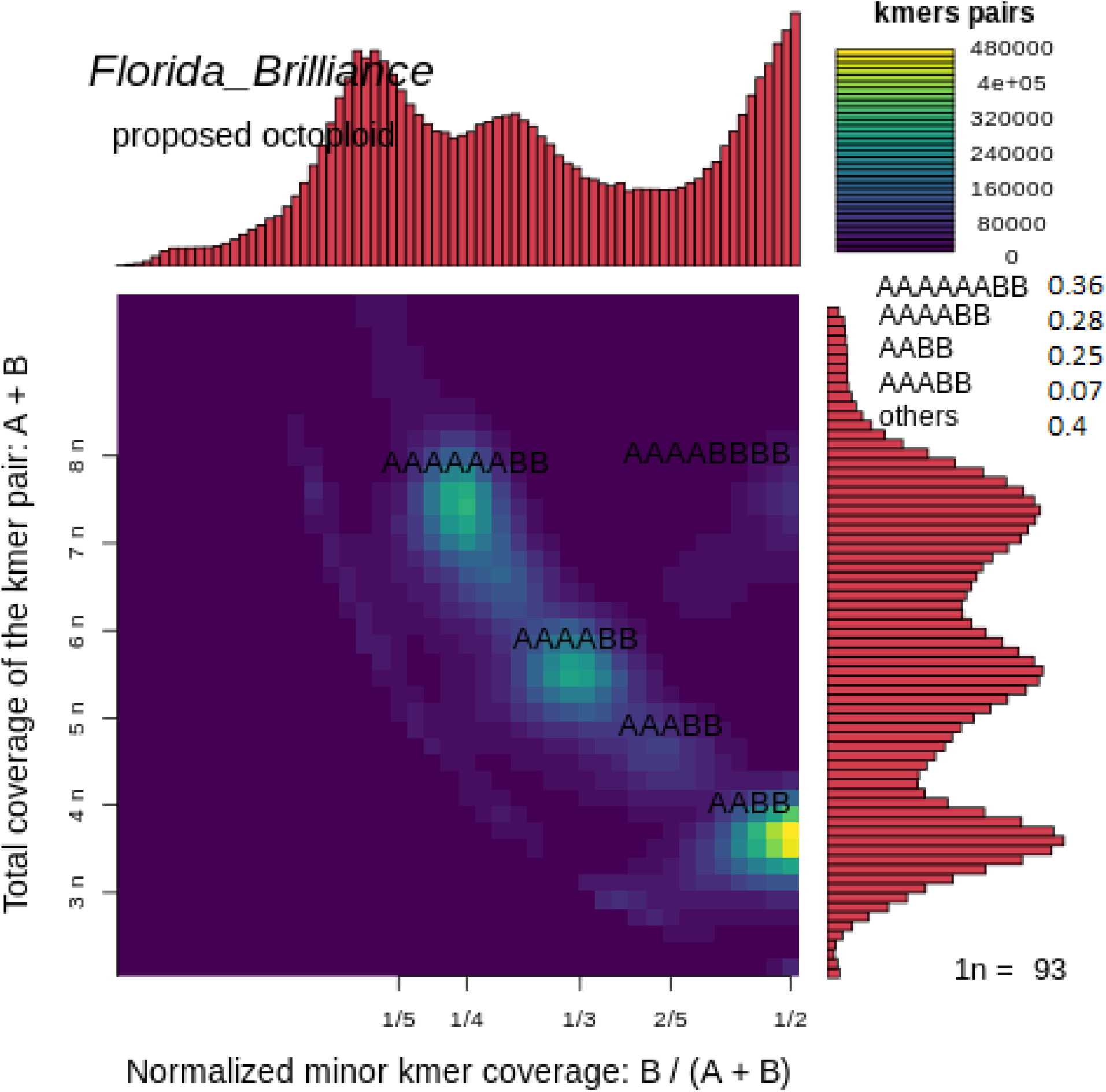
Smudgeplot for ‘Florida Brilliance’ (https://github.com/KamilSJaron/smudgeplot/). The brightness of each smudge is determined by the number of k-mer pairs, which was calculated by Jellyfish (9). The coloration indicates the approximate number of k-mer pairs per bin. The brightest smudge ‘AAAAAABB’ (0.36) suggests ‘Florida Brilliance’ is allo-octoploid with closely related paralogs. Other smudges including ‘AAAABB’ (0.28), ‘AABB’ (0.25) reflect that A and B are more closely related.

Hi-C reads were re-analyzed to scaffold the phased-1 and phased-2 assembly. Among 286 M Hi-C read pairs, 92.6% of Hi-C R1 and 91.7% of Hi-C R2 reads were mapped to the assembled genome (Supplementary Table 2). Unmapped read pairs, low quality pairs, and pairs with singleton were excluded, with 44.7% of uniquely mapped read pairs used for scaffolding using SALSA2. The unique Hi-C read pairs were visualized on the Hi-C contact map, showing 28 pairs of chromosomes on phase-1 and phased-2 genome assemblies (Figure 2).

**Figure 2.**
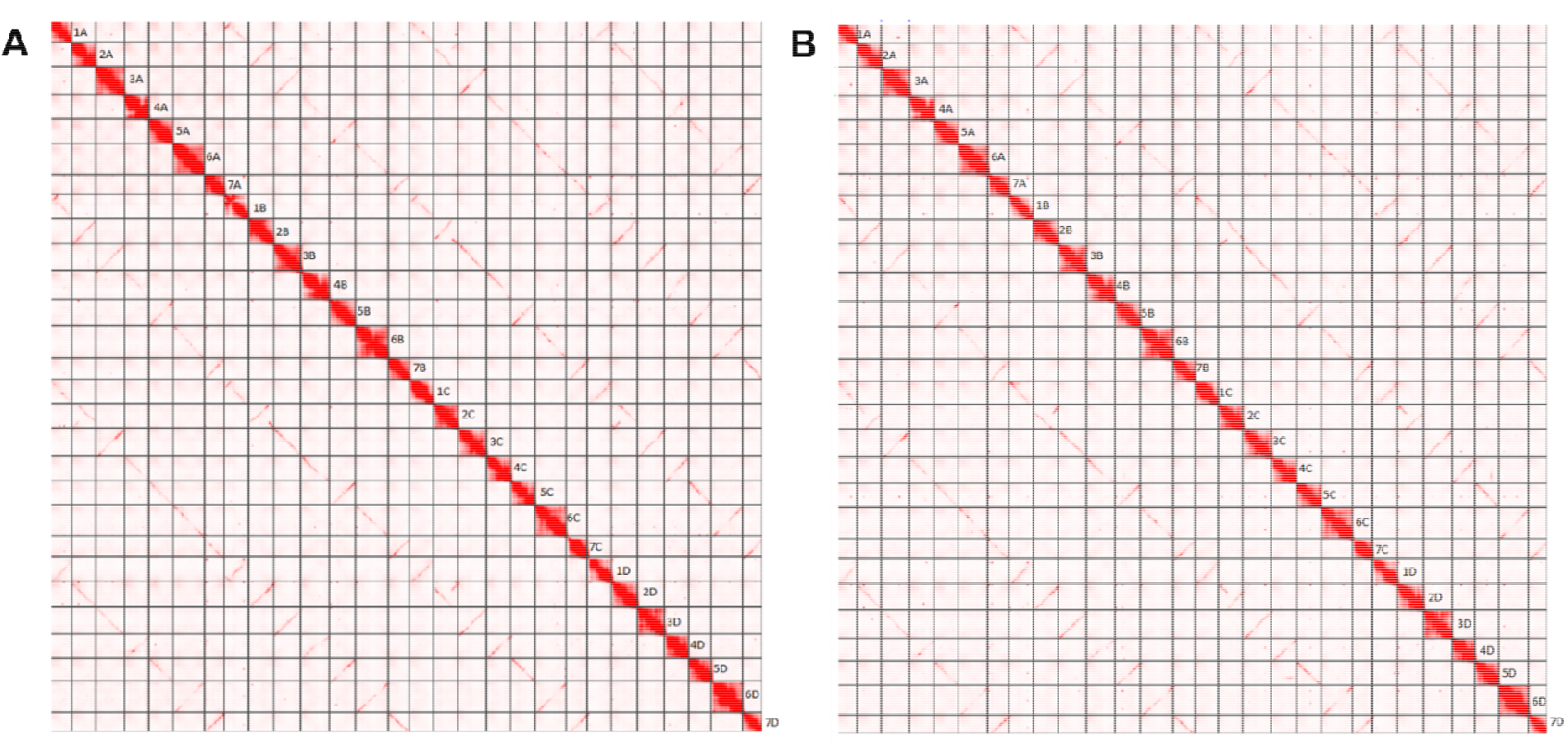
Hi-C contact map of phased-1 (A) and phased-2 (B) genome assemblies of ‘Florida Brilliance’. Each red pixel represents the Hi-C pairs. The dominant visual feature in every Hi-C heat map is the strong diagonal representing the Hi-C pairs of loci uniquely anchored in each haplotype-phased assembly.

Dense Hi-C read pairs were evenly distributed against whole assembly, indicating a low probability of mis-assemblies. Synteny between ‘Florida Brilliance’ and published reference genomes were visualized (Figure 3, Supplementary Figure 1). We first confirmed collinearity between phased-1 and phased-2 genome assemblies of ‘Florida Brilliance’ (Figure 3A). Alignments of the phased-2 assembly against the diploid *F*. *vesca* v4.0 showed a high degree of collinearity with the exception of major translocations on 1A and 2C (Figure 3B). In particular, ‘Florida Brilliance’ sub-genome A showed higher sequence similarity with *F*. *vesca* than other sub-genomes, which agrees with Hardigan et al. (5). Translocations on 3C, 4A, 4B and 4C were apparent when comparing with the genome of *F*. *chiloensis* (Figure 3D) (10). The genetic positions from a ‘Florida Brilliance’ linkage map and physical coordinates of SNPs showed good collinearity (Figure 4). Alignments of ‘Florida Brilliance’ phased-2 assembly against FaRR1 also displayed a high degree of collinearity (Figure 3C). Based on this alignment we applied the FaRR1 chromosome nomenclature reflecting the putative diploid origins of each respective sub-genome (A, B, C, D) (11). From each of the 56 pseudo-chromosomes of the combined phased −1 and −2 assemblies, we selected the most continuous pseudomolecule for the final FaFB2 assembly.

**Figure 3.**
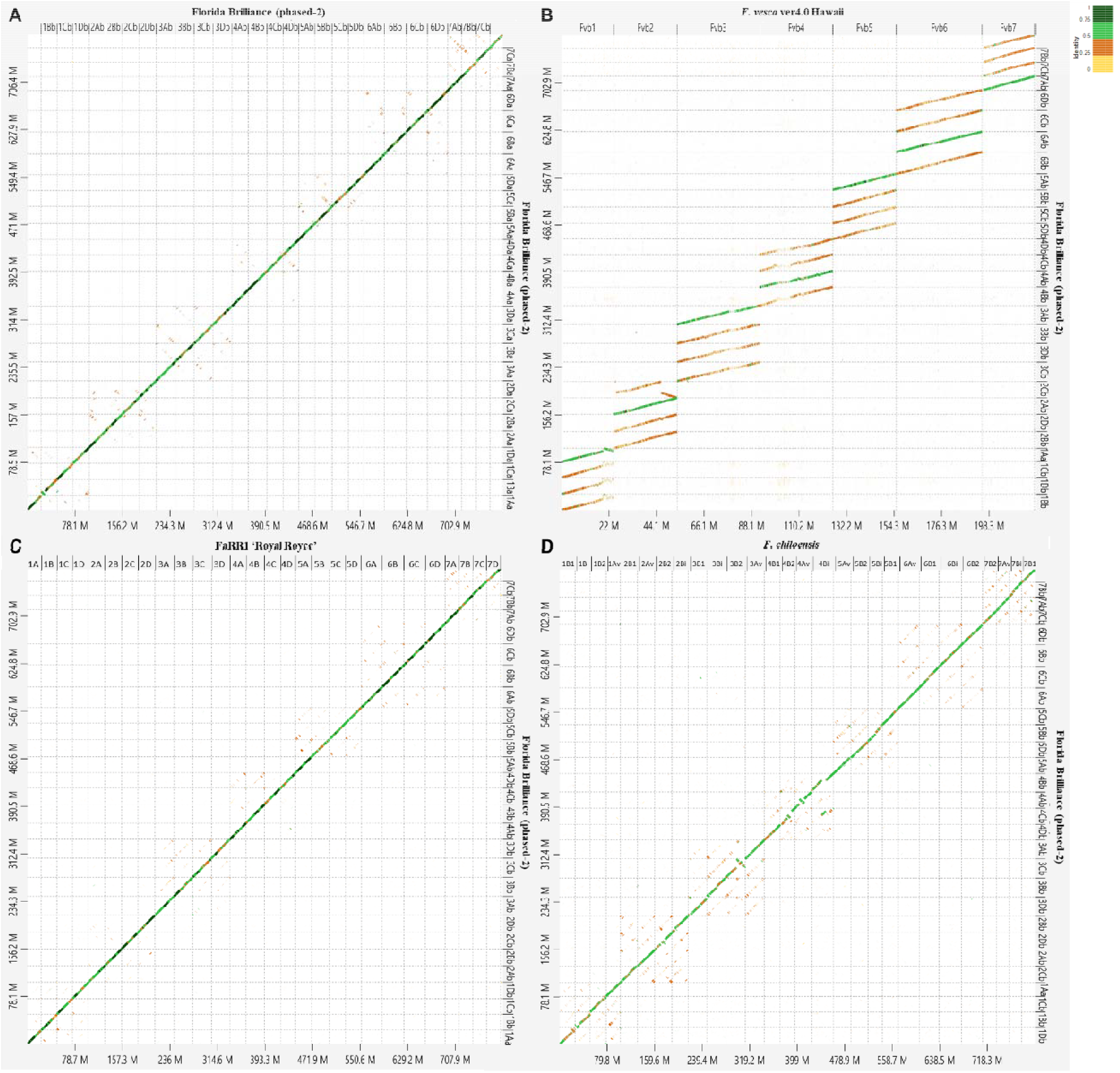
Dotplot of ‘Florida Brilliance’ phased-2 assembly to ‘Florida Brilliance’ phased-1 assembly (A), diploid *F*. *vesca* ver 4.0 (B)*, F*. × *ananassa* cv. Royal Royce (C), and *F*. *chiloensis* (D). Dot plots were produced using the DGENIE software and alignments with minimap2.

**Figure 4.**
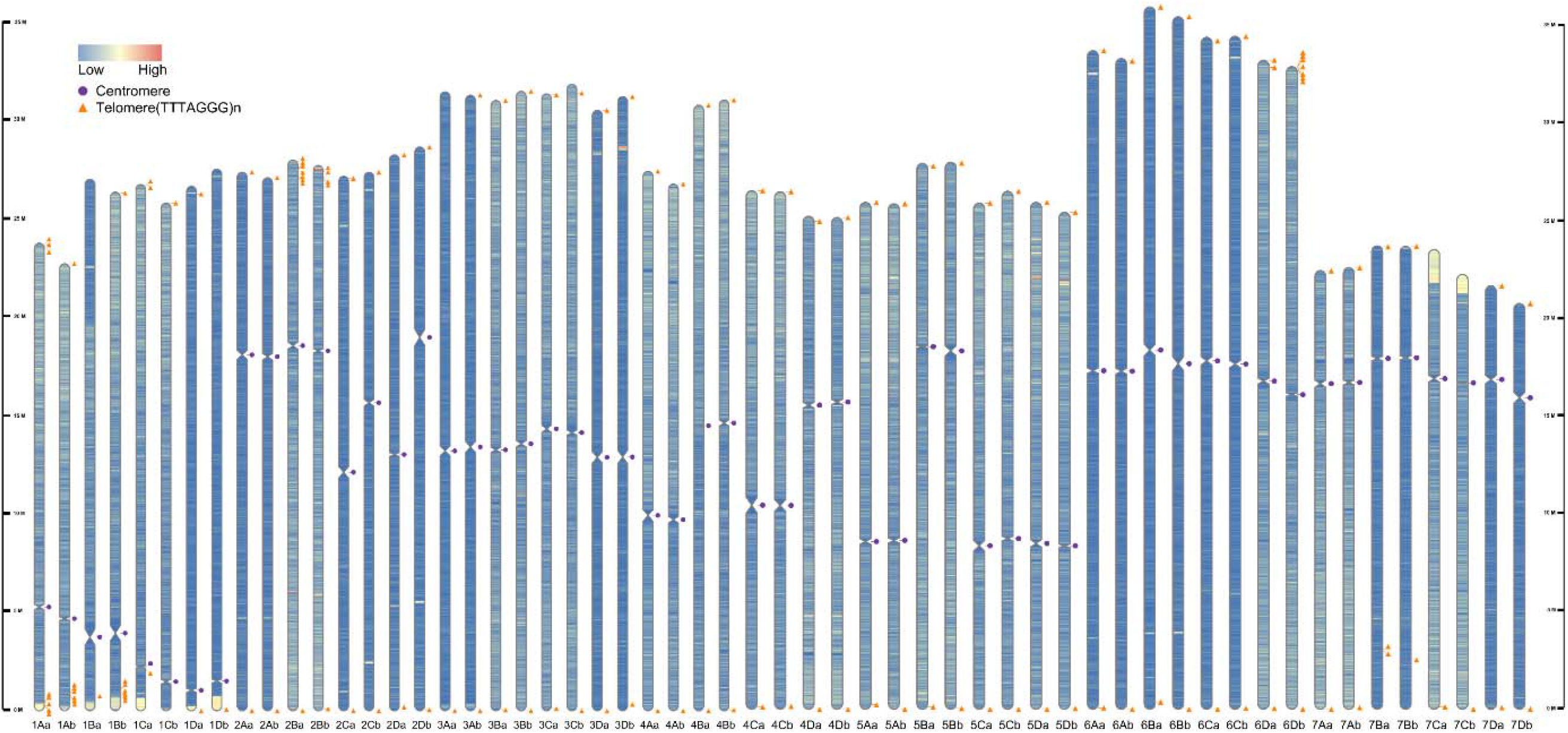
Telomere-to-telomere haplotype-phased genome assembly for octoploid strawberry ‘Florida Brilliance’. Triangle (Orange) represent the telomere sequences (5’-TTTAGGG-3’). Multiple triangles represent interstitial telomere-like sequence. Circles (purple) indicate the centromere region with low gene density and high density of repetitive sequences including LTR RTs and mini-satellites.

Telomeric motifs (5’-TTTAGGG-3’) were enriched in the terminal parts of the pseudo-chromosomes, allowing the identification of 103 telomeres (Figure 5). Putative telomeric sequences were found at the 5’ and/or 3’ ends in all 56 pseudo-chromosomes, and 50 were assembled telomere-to-telomere (Figure 5). Two pseudomolecules were potentially telomere-to-telomere, but the putative telomeric sequences were located 6 Mb or 7 Mb from end of chromosome 7A. Seven chromosomes had a short interstitial telomere-like sequence (Figure 5). Gene distribution along the chromosomes followed the typical distribution of monocentric plant genomes, and the positions of centromeres were similar between the phased-1 and phased-2 assemblies (Figure 5). We confirmed that the pericentric regions showed a high density of repeated sequences (Supplementary Figure 4). To evaluate the assemblies based on core eudicot genes, the genome quality was assessed using LTR assembly Index (LAI) (12). LAI for each chromosome ranged from 17.17 to 20.13. LAI for the combined phased-1 and phased-2 assemblies was 19.72 (Supplementary Table 3).

### Transcriptome Assembly and Gene Annotation

In total, 41.66% of the combined phased-1 and phased-2 assemblies were annotated as repetitive sequence. The majority of this repeat sequence was contributed by LTR transposable elements (Supplementary Table 4). Among them, 223,060 simple sequence repeats (SSRs) representing 0.16% of whole genome sequence were found (Supplementary Table 5). We obtained a redundant set of 123,319 transcripts by aligning Iso-Seq datasets to the combined phased-1 and phased-2 assemblies (supplementary file 1). Among redundant transcripts, 23,090 were generated from ‘Florida Brilliance’ Iso-seq and 94,229 were from ‘Royal Royce’ Iso-seq. After filtering, 83,070 non-redundant transcripts were retained and provided for gene annotation. BUSCO analysis of the transcript assemblies revealed 2,226 complete core eudicots genes (95.8%, 7.1% single-copy, 88.7% duplicated) with 1.0% fragmented and 3.2% missing core eudicots genes (Supplementary Table 6). Custom repeat libraries and non-redundant transcripts were provided for gene annotation with GenSAS v6.0 (13). Integrating ab initio predictions and evidence-based prediction, 196,403 genes remained in the final annotation with 98,384 models in the phase-1 assembly, 98,019 models in the phase-2 assembly (Supplementary Table 7), and 98,206 in the FaFB2 assembly (Supplementary Table 8). When classifying all predicted genes by sub-genome, sub-genome A possessed 27.4% of gene content (Tables S7 and S8).

## DISCUSSION

Our smudgeplot displayed a genome structure of cultivated strawberry based on K-mer analysis that supports genomic structure of ‘AAAAAABB’, similar to ‘AvAvBiBiB1B1B2B2’ (3). When examined by sub-genome, we observed sub-genome dominance: A > B > C > D based on the proportion of complete vs fragmented or missing genes in the ‘Florida Brilliance’ genome assembly, as with ‘Royal Royce’(5), and *F*. *chiloensis* (10). Genome structure ‘AAAAAABB’ along with sub-genome dominance might support the model hypothesizing one subgenome arising from cytoplasm donor *F*. *vesca*, one subgenome from diploid *F*. *iinumae*, and two from an unknown diploid close to *F*. *iinumae* (2).

Since the first report of a chromosome-scale reference genome by Edger et al (2), several octoploid strawberry (*F*. × *ananassa*) reference genomes using HiFi reads have been generated (2, 5, 14, 15). The high accuracy of HiFi reads notably improved the assembly quality for octoploid strawberry. Interestingly, the N50 of ‘Florida Brilliance’ initial phased-1 and phased-2 (23.7 and 26.7 Mb) assemblies were higher than other recent genomes: Wongyo3115 (9.84 Mb), ‘Royal Royce’ (11 Mb), *F*. *chiloensis* (10.8 Mb), and ‘Reikou’ (3.9 Mb). Multiple contigs with lengths similar to those after scaffolding indicate multiple single-contig chromosome assemblies (Supplementary Figure 2). The final haploid assemblies of 784.9 Mb and 781.0 Mb for phased-1 and phased-2 are in accordance with assembly length of ‘Royal Royce’ (784 Mb) (5). When comparing the corresponding chromosome lengths between ‘Florida Brilliance’, ‘Royal Royce’ and ‘Camarosa’, we confirmed a high similarity between ‘Royal Royce’ and ‘Florida Brilliance’ (Supplementary Figure 3). This indicates that haplotype-phased genome assemblies of a highly heterozygous allo-octoploid can be successfully generated without parental reads (6).

Telomeres, basic structure of eukaryotic chromosomes, are typically tandemly arranged mini-satellites following the formula (TxAyGz)n at both ends of chromosomes (16). Telomere-to-telomere genome assemblies are needed to ensure that all gene content is accounted for. Recently, putative telomeric sequences (5’-TTTAGGG-3’) of octoploid strawberry were found at both ends in 7 of the 28 pseudo-chromosomes of *F*. *chiloensis* (10). In the present study, 90% of putative telomeres were found in combined phased-1 and phased-2 genome assemblies for ‘Florida Brilliance’ (Figure 1) and were indicative of a high-quality assembly.

Average LAI scores for all 56 pseudo-chromosomes were 19.72, and were comparable to those of ‘Royal Royce’ (18.33-19.22) and *F*. *vesca* Hawaii 4 (18.44) (5). Genome assemblies based on NGS technologies have typically possessed very low sequence continuity compared to Sanger-based technique or BAC-based scaffoldings (12). The BUSCO score of ‘Florida Brilliance’ (99.2%) is highest among published reference octoploid strawberry genomes, when compared to ‘Camarosa’ (96.2%), ‘Royal Royce’ (98.1%), ‘Wongyo 3115’ (94.1%), and *F*. *chiloensis* (99.1%), indicating a high level of completeness. In the combined phased-1 and phased-2 genome assembly, there was 652 Mb of repetitive sequence accounting for 41.66% of the genome, which is higher than published references ‘Royal Royce’ (38.4%), ‘Camarosa’ (36.0%), and ‘Wongyo 3115’ (38.8%).

The total gene number of FaFB2 was 104,099 (Supplementary Table 8), similar to ‘Royal Royce’ (101,721) (5) and ‘Camarosa’ (108,087) (2) but different from ‘Wongyo 3115’ (151,892) (14) and ‘Reikou’ (167,721) (15). The different number of genes among published genomes could have resulted from the difference in the implementation of default parameters for the gene prediction. Although the total number of genes differs among the genomes available, sub-genome A representing diploid progenitor *F*. *vesca* occupied the largest proportion (27.5%) (Tables 8), similar to ‘Royal Royce’ (27%) (5). We also confirmed sub-genome dominance (A > B > C > D) based on the total number of genes estimated for each sub-genome (Tables S7 and S8).

Several chromosomal rearrangements in the *F*. *chiloensis* reference relative to *F*. *vesca* and ‘Royal Royce’ (*F*. × *ananassa*) have been reported (10) and are also visible when compared to ‘Florida Brilliance’ (Figure 3). For example, the alignment against *F*. *chiloensis* shows disagreement in continuity in 3 chromosomes (Fchil3-B2, Fchil4-Av, and Fchil4-Bi) (Figure 3D). The chromosomal rearrangements were not found in 3 chromosomes between ‘Florida Brilliance’ and ‘Royal Royce’. However, further investigation is necessary to resolve whether these disagreements are misassembles or true rearrangement.

## CONCLUSIONS

Our study demonstrates the assembly of a highly complex polyploid genome with a combination of PacBio HiFi long-read sequencing and Hi-C data. This strategy led to the completion of the first telomere-to-telomere, haplotype-phased assembly of an octoploid strawberry, ‘Florida Brilliance’. This high-quality assembly was generated without parental sequence data and was assigned sub-genomes A, B, C, and D representing sequence similarity with *F*. *vesca* and having high collinearity with the octoploid reference FaRR1 (‘Royal Royce’). This newly-available FaFB2 assembly will contribute to the discovery of genes for important traits and accelerate the development of breeding tools and approaches.

## MATERIALS AND METHODS

### Sample collection and sequencing

Young leaf tissue was covered with black plastic bags and kept in a greenhouse for 2 weeks. The etiolated leaf tissue was harvested for the DNA extraction. Leaves were frozen then submitted to DNA Link (Seoul, South Korea) for genomic DNA extraction and library preparation. The single-molecule real-time sequencing (SMRT) bell library was constructed using a PacBio DNA Template Prep Kit 1.0 (Pacific Biosciences). Quality and quantity of each library was checked using a 2100 Bioanalyzer (Agilent Technologies). The SMRT Bell-Polymerase complex was constructed using a PacBio Binding Kit 2.0 (Pacific Biosciences) based on the manufacturer’s instructions. The complex was loaded onto five SMRT cells (Pacific Biosciences, Sequel SMRT Cell 1M v2) and sequenced using Sequel Sequencing Kit 2.1 (Pacific Biosciences, Sequel SMRT Cell 1M v2). For each SMRT cell, 1 × 600 min movies were captured using the Sequel sequencing platform (Pacific Biosciences) at DNA Link (Seoul, South Korea). The quality of Hifi was measured with LongQC (17).

For transcriptome sequencing, six ‘Florida Brilliance’ tissues were used for construction of PacBio Iso-Seq libraries. All ‘Florida Brilliance’ tissues were harvested from experimental fields at the Gulf Coast Research and Education Center (GCREC) in Wimauma, Florida between 14:00 and 16:00 in February 2021. Roughly 10 grams of 6 tissues were collected from 5 plants: flowers, green fruit, red fruit, crown, leaf, and root. All tissues were flash frozen at −80□ after washing with water. Total RNA was extracted using Spectrum™ Plant Total RNA Kit (Sigma-Aldrich, St. Louis, MO, United States) according to manufacturer’s instructions. To remove any traces of DNA removed with DNAse I (Invitrogen) and re-suspended in a total volume of 50 μl of RNase-free water. The six ‘Florida Brilliance’ RNA samples were submitted to DNA Link for preparation of Iso-Seq library. Pooled libraries were sequenced on SMRT-cells on the Sequel I platform.

### *De Novo* genome assembly and validation

Overall genome characteristics including genome size and repetitive elements were estimated using PacBio HiFi data by K-mer spectrum distribution analysis for *k□* = □21 in KMC3(18) and GENOMESCOPE v 2.0 (19). The HiFi and Hi-C reads were used to produce a haplotype-phased assembly without sequencing of parents using Hifiasm (6). Hifiasm was run with the following command according to developer’s recommendation for heterozygous crops: hifiasm -o <outputPrefix> -t <nThreads> -D10 <Hifi-reads.fasta> --h1 <Hi-C_reads1> --h2 <Hi-C_reads2>. SALSA2, which is Hi-C-based scaffolding programs, was used for scaffolding contigs. Hi-C Reads were mapped by Arima-HiC mapping pipeline (https://github.com/ArimaGenomics/mapping_pipeline). After mapping Hi-C reads to each phased genome assembly using BWA v0.7.17 (20) and sequence alignment map (SAM) format was converted to bed format using SAMtools (21) and BEDTools (22) prior to SALSA2 scaffolding (https://github.com/marbl/SALSA; (23), which was run with parameters -e GATC -m yes. Hi-C reads were mapped to chromosomes using HiC-Pro (24) in order to assess the quality of assembly. The interaction matrix of whole chromosomes were visualized with heatmaps. Remaining contigs were scaffolded and oriented based on ‘15.89-25’ (https://www.rosaceae.org) reference genome using Ragtag (25).

### Scaffold validation using a genetic linkage map

A high-density genetic map was developed using a total of 169 F1 individuals from a cross between ‘Florida Brilliance’ and breeding selection FL 16.33-8. Axiom™ IStraw35 SNP arrays were used to genotype all 169 F1 individuals. Markers were filtered to have <5% missing data and fit segregation ratios of 1:1 and 1:2:1 (α = 0.05). Marker genotype calls were recoded to fit the Joinmap 4.1 linkage mapping requirement. For example, markers with paternal segregation (AA × AB or BB × AB) coded as “nn × np”; markers with maternal segregation (AB × AA or AB × BB) coded as “lm × ll”: and markers segregating in both parents (AB × AB) coded as “hk × hk.” Mapping was conducted in an iterative process using the maximum likelihood algorithm in JoinMap 4.1 with default settings. After each round of mapping, a graphical genotyping approach was applied to identify singletons to fix the marker order and regions with low marker density or gaps caused by segregation distortion. A final linkage map of ‘Florida Brilliance’ consisting of 10,269 SNP markers was used to validate the scaffolds from the FaFB2 whole genome assembly. SNP probes sequences used in the construction of linkage maps were mapped to the FaFB2 assembly sequence using blastn procedure (26). Alignments were filtered to retain markers if they matched to unique sequence position in the FaFB2 phased genome assembly and with a maximum of 2 mismatches in the second-best hit. The alignments were queried to detect potentially problematic scaffolds mapped with SNP probes from different LGs. The number of scaffolds with SNP probes mapped from different LGs was used as a metric in the quality assessment of FaFB2 assembly.

### Assembly quality evaluation

Genome assembly statistics were calculated using QUAST version 5.0.266. Merqury version 1.3 were used to measure assembly consensus quality value (QV), evaluates assembly based on efficient K-mer set operations (27). The completeness of the haploid assemblies and protein-coding gene annotations were assessed with the BUSCO database (28). The scaffolds were inspected on the Hi-C contact map. Hi-C reads were trimmed with Homer (29) and mapped to the both haploid assembly using HiC-Pro version 3.0 (24) and visualized in Juicebox version 1.11 (30). LAI (12) for each sub-genome was calculated using LTR-retriever (31) along with whole-genome TE-annotations and intact LTR retrotransposons identified by EDTA (32).

### Repetitive annotation

Transposable elements (TEs) were annotated using EDTA v1.9.6 with default parameters (33). The TE annotation library was generated by EDTA in a separate run. TE regions of both haploid assemblies were masked by ReapeatMasker v4.1.1 provided with the repeat library. Simple sequence repeats (SSRs) or microsatellites were mined using SSR Finder (13) on Genome Sequence Annotation Server v6.0 (GenSAS;https://www.gensas.org). Telomeric repeats were annotated using BIOSERF (34).

### Transcriptome assembly and gene annotation

To increase the accuracy of gene annotation, we generated a transcriptome assembly containing possible sets of transcripts from ‘Florida Brilliance’ and *F*. × *ananassa* expression data publicly available (Supplementary Table 9). Octoploid strawberry ‘Royal Royce’ Iso-Seq reads were trimmed using Trimmomatic version 0.39 and mapped to two phased-assemblies using HISAT v2.2.1 (35) with default parameters. The ‘Florida Brilliance’ and ‘Royal Royce’ Iso-Seq reads were aligned to the assemblies using minimap v2.2.1 (36). Reads alignment were converted to Binary alignmented map (BAM) format with SAMTools. Reference-guided transcriptome assembly was performed using StringTie v2.1.4 (37) with the Iso-Seq alignments as input. StringTie2 was run with default parameters and with the addition of long read (-L) mode. Mikado v2 (38) was used to generate a non-redundant set of transcript assemblies with best-scoring transcript evidence at each locus. Match scores were measured for all transcriptome assemblies against the UniProt protein database using BLASTX. TransDecoder v5.5.0 (https://github.com/TransDecoder/TransDecoder) were used to predict the best six-frame translations of the transcriptome assemblies from StringTie2, then splice junctions for all merged RNA alignments were predicted with Portcullis v1.2.2 (39). The Mikado scoring for any transcript assemblies derived from Iso-Seq alignments was modified over RNA-Seq alignments. TransDecoder was used to filter non-redundant, polished transcripts generated by Mikado to obtain best ORF scores. The ‘Florida Brilliance’ genome assembly was annotated using GenSAS v6.0 (13). Non-redundant transcript assemblies derived from Mikado2 pipeline were provided as EST evidence. Transcripts were aligned to combined phased-1 and phased-2 assemblies using BlastN (26). The alignments were combined with results of gene prediction using AUGUSTUS v3.1.1 (40) to generate an official gene set and identify predicted transcripts within each masked assembly (EVidenceModeler) (41, 42). Function of predicted transcripts were annotated based on alignment using BlastP v2.2.28 to the UniProtKB database (43).

### Collinearity and synteny

Proteins of diploid strawberry progenitor *F*. *vesca* were collected for all-against-all alignments to predicted proteins for Octoploid strawberry ‘Florida Brilliance’. These alignments were passed to MCScanX to identify synteny blocks (44). DNA level synteny between *F*. *vesca*, *F*. × *ananassa*, *F*. *chiloensis* and two phased genome assemblies of ‘Florida Brilliance’ were all plotted using D-GENIES (45) with default parameters after aligning with minimap2.

### Availability of Supporting Data and Materials

The chromosome-level goose genome assembly, annotation files, and other supporting data are available via the Genome Database for Rosaceae (GDR) (13) and DRYAD database. The source of Iso-seq raw data used are listed in deposited in the DRYAD database ().

## SUPPLEMENTARY TABLES

**Supplementary Table 1.** Statistics of HiFi reads (QV ≥ 20) generated from iso-seq sequences from 6 tissues of ‘Florida Brilliance’

| Tissue | Yield (bp) | Number of reads | Average length |
| --- | --- | --- | --- |
| Leaf | 125,573,371 | 50,973 | 2,463.5 |
| Crown | 138,458,257 | 56,707 | 2,441.6 |
| Flower | 120,850,566 | 51,203 | 2,360.2 |
| Green fruit | 132,446,690 | 54,049 | 2,450.5 |
| Red fruit | 132,761,812 | 49,638 | 2,674.6 |
| Root | 116,103,170 | 45,577 | 2,547.4 |
| Total | 766,193,866 | 308,147 |  |
| Average |  |  | 2,486.5 |

**Supplementary Table 2.**
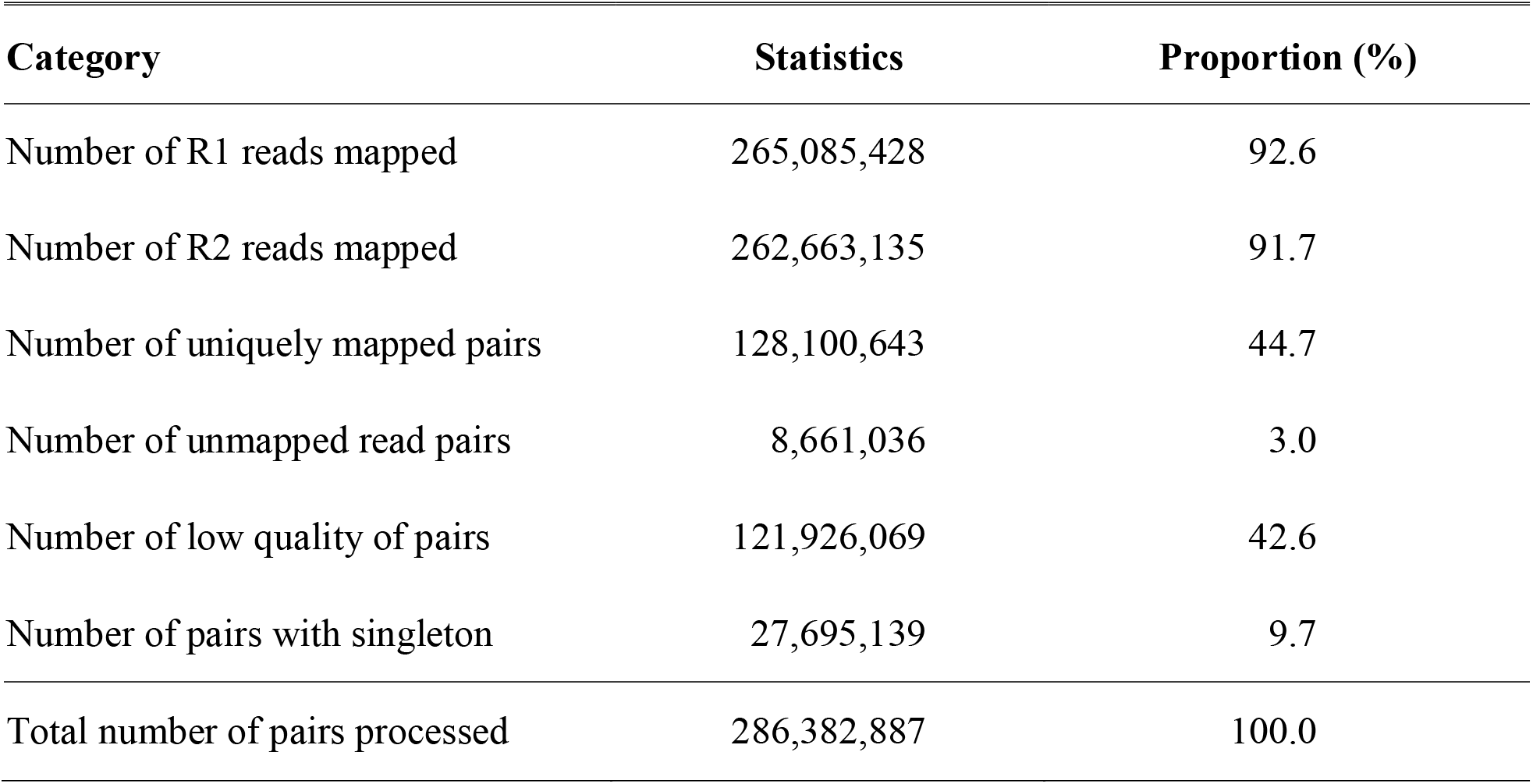
Statistics of genome assembly and Hi-C analysis for octoploid strawberry ‘Florida Brilliance’.

**Supplementary Table 3.** LTR Assembly Index (LAI) scores for ‘Florida Brilliance’ phase-1 and phase-2 subgenomes.

| **Genome assembly** | **Sub-genome** | **Haplotype phase** | **LAI** |
| --- | --- | --- | --- |
| Florida Brilliance | A | Haplotype-1 | 19.72 |
| Florida Brilliance | A | Haplotype-2 | 19.27 |
| Florida Brilliance | B | Haplotype-1 | 17.17 |
| Florida Brilliance | B | Haplotype-2 | 18.44 |
| Florida Brilliance | C | Haplotype-1 | 18.83 |
| Florida Brilliance | C | Haplotype-2 | 19.17 |
| Florida Brilliance | D | Haplotype-1 | 19.75 |
| Florida Brilliance | D | Haplotype-2 | 20.13 |
| **Total** |  |  | **19.72** |

**Supplementary Table 4.** Classification and distribution of repetitive DNA elements identified in the genome assembly for ‘Florida Brilliance’ by EDTA pipeline.

|  | Class | Sub-Class | Count | Length of masked sequence (bp) | Proportion of masked sequence (%) |
| --- | --- | --- | --- | --- | --- |
| LTR | Class I |  |  |  |  |
|  |  | Copia | 99,547 | 73,506,892 | 4.69 |
|  |  | Gypsy | 174,983 | 184,244,965 | 11.77 |
|  |  | unknown | 286,120 | 149,607,484 | 9.55 |
| TIR | Class II |  |  |  |  |
|  |  | CACTA | 148,100 | 59,217,274 | 3.78 |
|  |  | Mutator | 215,008 | 67,488,697 | 4.31 |
|  |  | PIF_Harbinger | 55,008 | 20,728,437 | 1.32 |
|  |  | Tc1_Mariner | 4,971 | 1,709,712 | 0.11 |
|  |  | hAT | 84,543 | 33,197,559 | 2.12 |
| NonTIR | Class II |  |  |  |  |
|  |  | helitron | 142,472 | 62,596,795 | 4.00 |
| **Total** | **-** | **-** | **1,210,752** | **652,297,815** | **41.66** |

**Supplementary Table 5.** Simple sequence repeats of octoploid strawberry genome assembly for ‘Florida Brilliance’ using SSR Finder.

| **SSR type** | **Frequency (#)** | **Proportion (%)** | **Size (bp)** | **Genome coverage<sup>z</sup>**<br>**(%)** |
| --- | --- | --- | --- | --- |
| Dinucleotide | 80,951 | 36.29 | 820,359 | 0.05 |
| Trinucleotide | 52,060 | 23.34 | 593,417 | 0.04 |
| Tetranucleotide | 65,389 | 29.31 | 737,447 | 0.05 |
| Pentanucleotide | 15,126 | 6.78 | 217,899 | 0.01 |
| Hexanucleotide | 9,533 | 4.27 | 164,112 | 0.01 |
| **Total** | **223,060** | **100.00** | **2,533,234** | **0.16** |

**Supplementary Table 6.** Benchmarking universal single-copy orthologs (BUSCO) analysis of 123,319 transcriptome assemblies for ‘Florida Brilliance’.

|  | embryophyta_odb10 | eudicots_odb10 |
| --- | --- | --- |
| Complete BUSCOs (C) | 1,556 (96.4%) | 2,226 (95.8%) |
| Complete and single-copy BUSCOs (S) | 108 (6.7%) | 164 (7.1%) |
| Complete and duplicated BUSCOs (D) | 1,448 (96.4%) | 2,062 (88.7%) |
| Fragmented BUSCOs (F) | 21 (1.3%) | 23 (1.0%) |
| Missing BUSCOs (M) | 37 (2.3%) | 77 (3.2%) |
| Total BUSCO groups searched | 1,614 (100.0%) | 2,326 (100.0%) |

**Supplementary Table 7.** Genes predicted in the combined phased-1 and phased-2 genome assembly of ‘Florida Brilliance’.

| **Chromosome** | **Sub-genome in phased-1 assembly** |  |  |  |  | **Sub-genome in phased-2 assembly** |  |  |  |  |
| --- | --- | --- | --- | --- | --- | --- | --- | --- | --- | --- |
|  | **A** | **B** | **C** | **D** | **Total** | **A** | **B** | **C** | **D** | **Total** |
| Chr01 | 3,258 | 2,879 | 2,955 | 2,751 | 11,843 | 3,151 | 2,917 | 2,717 | 2,922 | 11,707 |
| Chr02 | 3,992 | 3,693 | 3,382 | 3,531 | 14,598 | 3,941 | 3,659 | 3,361 | 3,511 | 14,472 |
| Chr03 | 4,306 | 3,692 | 3,686 | 3,420 | 15,104 | 4,251 | 3,774 | 3,724 | 3,468 | 15,217 |
| Chr04 | 3,527 | 3,203 | 3,008 | 2,874 | 12,612 | 3,482 | 3,185 | 3,095 | 2,904 | 12,666 |
| Chr05 | 3,667 | 3,500 | 3,077 | 3,174 | 13,418 | 3,667 | 3,535 | 3,136 | 3,109 | 13,447 |
| Chr06 | 5,053 | 4,522 | 4,352 | 4,216 | 18,143 | 5,017 | 4,493 | 4,365 | 4,221 | 18,096 |
| Chr07 | 3,400 | 3,017 | 3,466 | 2,783 | 12,666 | 3,371 | 3,074 | 3,187 | 2,782 | 12,414 |
| Total | 27,203 | 24,506 | 23,926 | 22,749 | 98,384 | 26,880 | 24,637 | 23,585 | 22,917 | 98,019 |
| Percentage (%) | 27.65 | 24.91 | 24.32 | 23.12 | 100 | 27.42 | 25.13 | 24.06 | 23.38 | 100 |

**Supplementary Table 8.** Genes predicted in the FaFB2 of ‘Florida Brilliance’.

| **Chromosome** | **Sub-genome** |  |  |  |  |
| --- | --- | --- | --- | --- | --- |
|  | **A** | **B** | **C** | **D** | **Total** |
| Chr01 | 3,258 | 2,917 | 2,717 | 2,751 | 11,643 |
| Chr02 | 3,992 | 3,659 | 3,382 | 3,531 | 14,564 |
| Chr03 | 4,251 | 3,774 | 3,686 | 3,420 | 15,131 |
| Chr04 | 3,527 | 3,203 | 3,008 | 2,874 | 12,612 |
| Chr05 | 3,667 | 3,535 | 3,136 | 3,109 | 13,447 |
| Chr06 | 5,053 | 4,522 | 4,352 | 4,216 | 18,143 |
| Chr07 | 3,400 | 3,017 | 3,466 | 2,783 | 12,666 |
| Total | 27,148 | 24,627 | 23,747 | 22,684 | 98,206 |
| Percentage (%) | 27.6 | 25.1 | 24.2 | 23.1 | 100.0 |

## SUPPLEMENTARY FIGURES

**Supplementary Figure 1.**
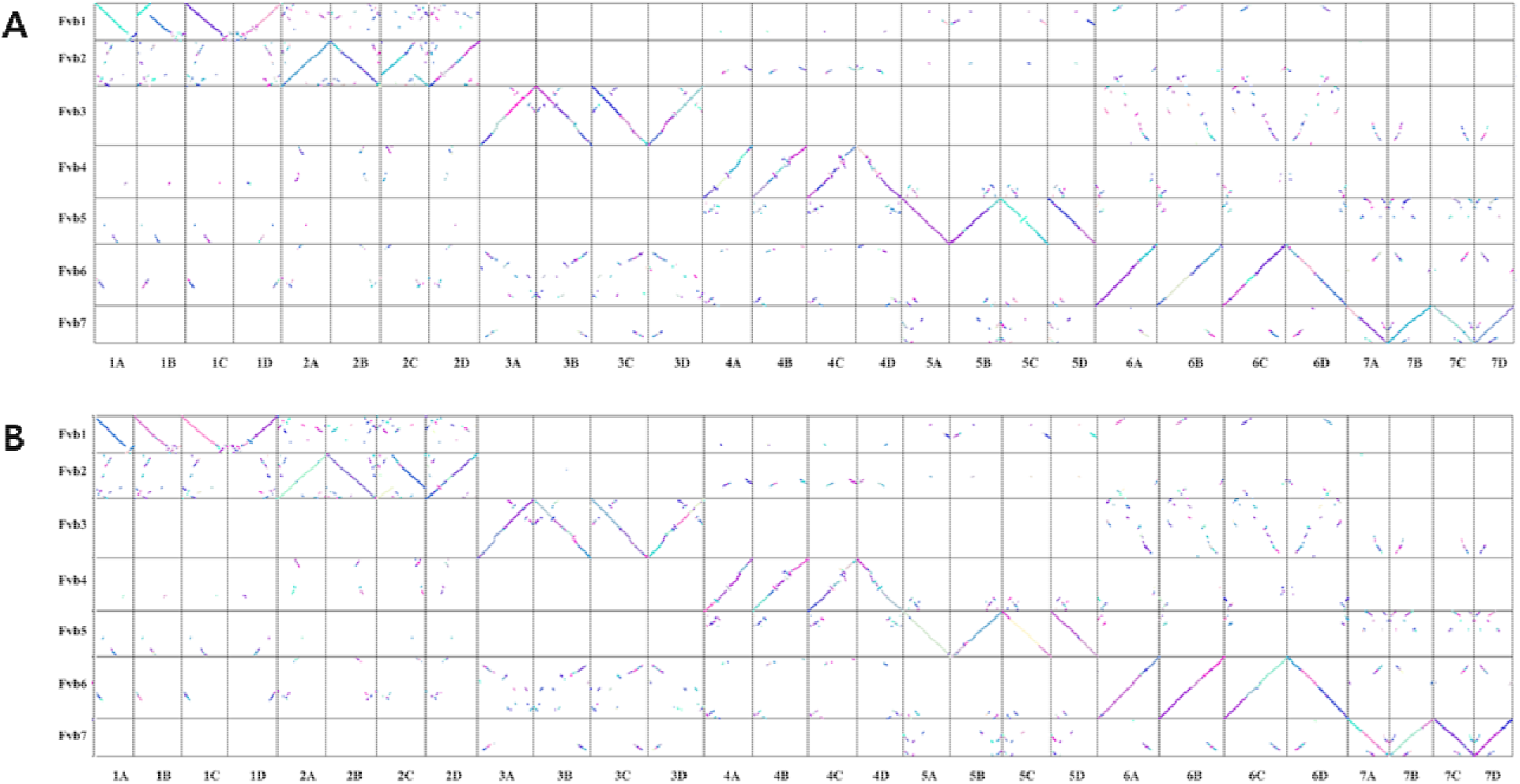
Dotplot of collinear gene pairs between ‘Florida Brilliance’ and *F*. *vesca*. Homologous genes of phased-1 (A) and phased-2 (B) assemblies were compared against diploid *F*. *vesca*.

**Supplementary Figure 2.**
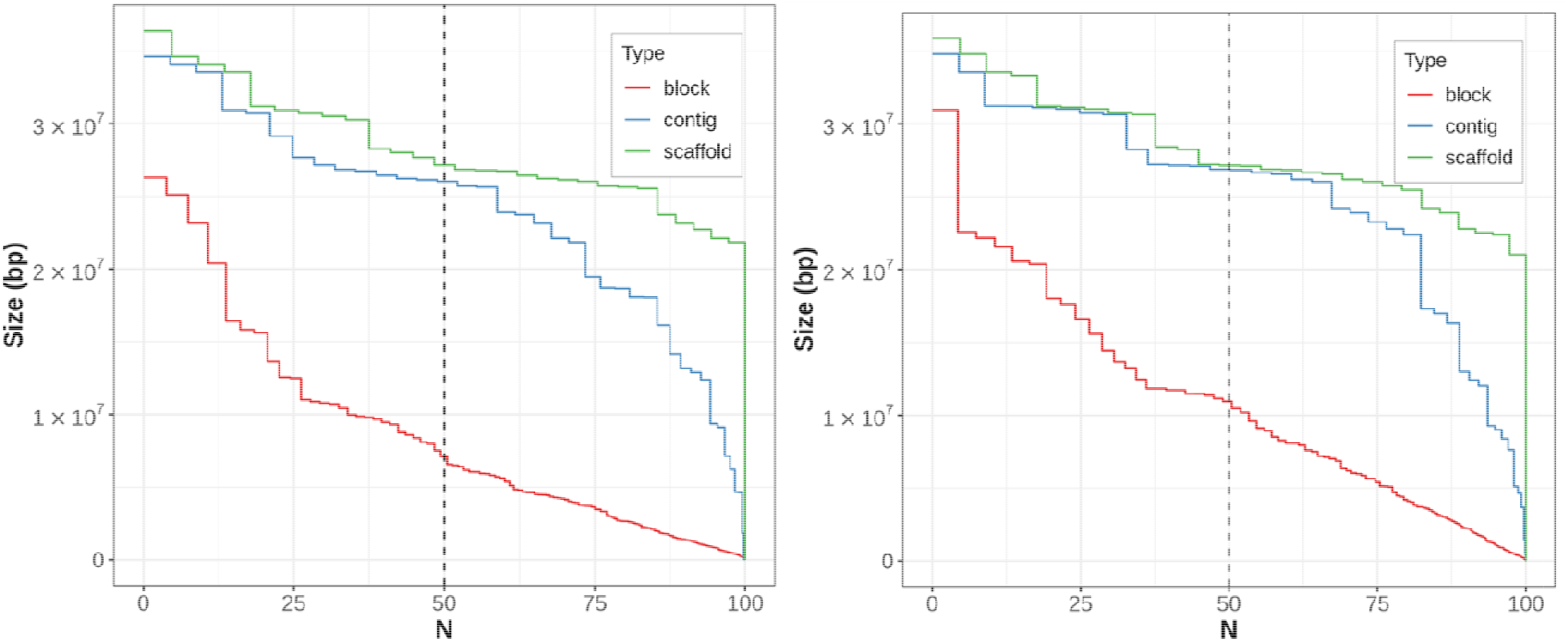
N50 of phased block, contig, and scaffolds in phased-1 (A) and phased-2 assembly (B).

**Supplementary Figure 3.**
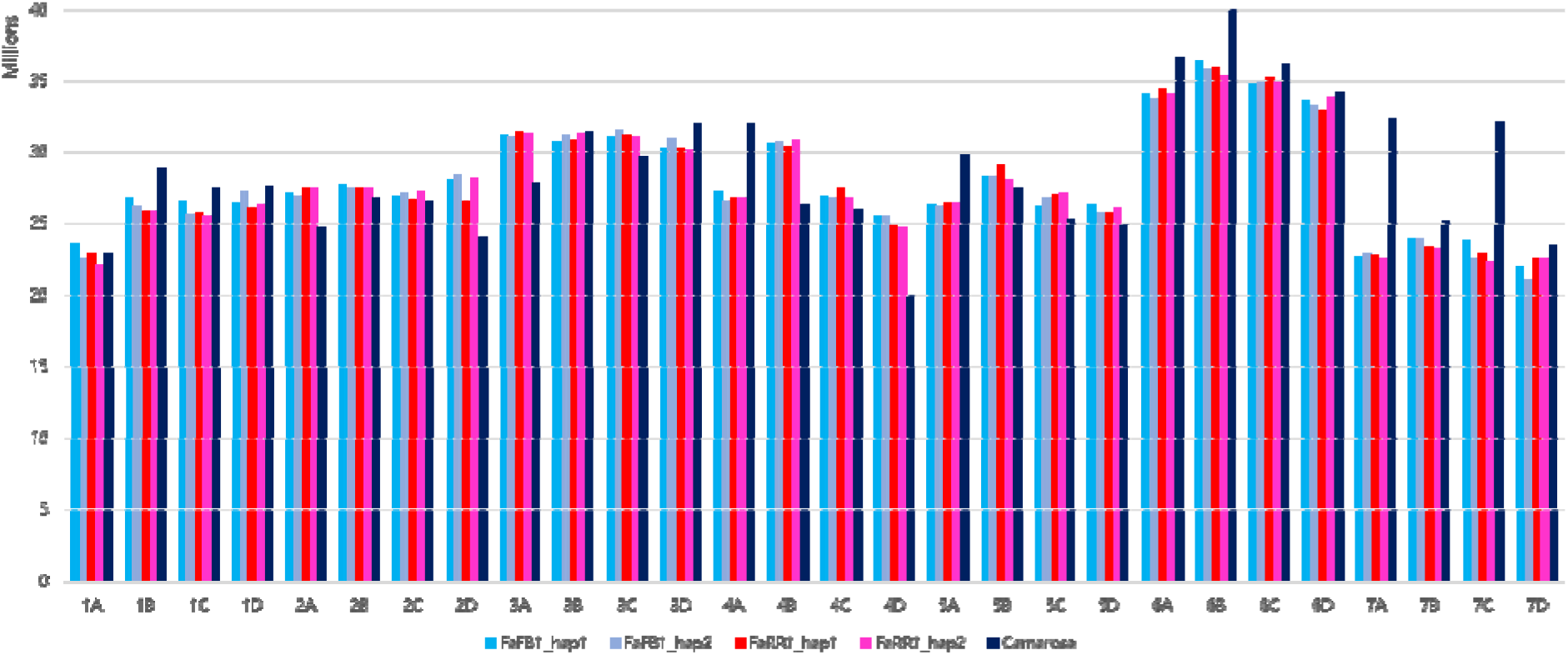
Chromosome length of ‘Florida Brilliance’, ‘Royal Royce’ and ‘Camarosa’ assemblies. ‘Florida Brilliance’ and ‘Royal Royce’ consist of two phased assemblies (phased-1 and phased-2).

**Supplementary Figure 4.**
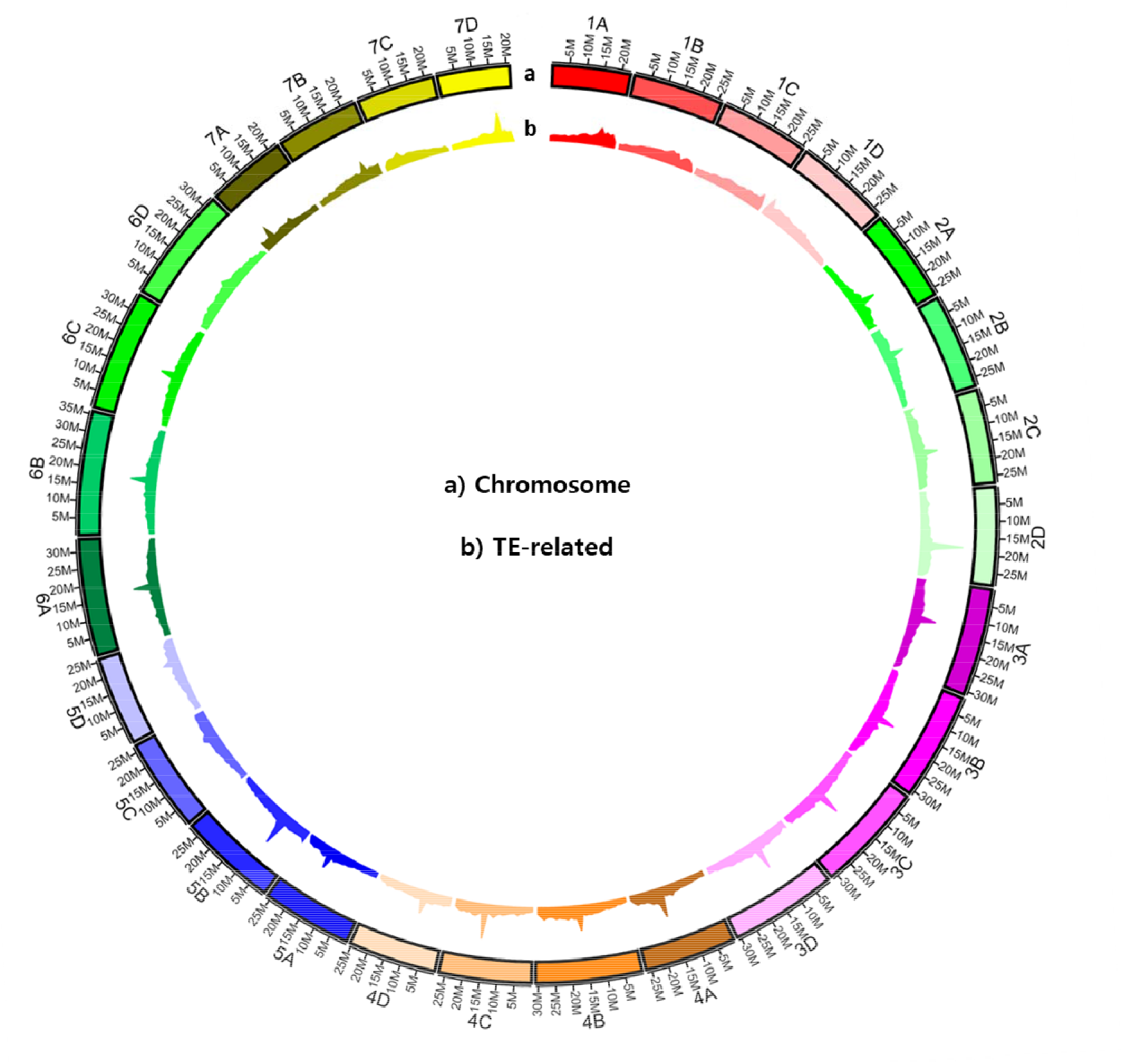
The genomic features of the octoploid strawberry genome. a, chromosome length. b, density of TE-related sequence (number of TE per Mb).

